# Revisiting the early evolution of Cyanobacteria with a new thylakoid-less and deeply diverged isolate from a hornwort

**DOI:** 10.1101/2021.02.18.431691

**Authors:** Nasim Rahmatpour, Duncan A. Hauser, Jessica M. Nelson, Pa Yu Chen, Juan Carlos Villarreal A., Ming-Yang Ho, Fay-Wei Li

**Author notes:** Authors of correspondence: Fay-Wei Li and Ming-Yang Ho.

## Abstract

Cyanobacteria have played pivotal roles in Earth’s geological history especially during the rise of atmospheric oxygen. However, our ability to infer the early transitions in Cyanobacteria evolution has been limited by their extremely lopsided tree of life—the vast majority of extant diversity belongs to Phycobacteria (or “crown Cyanobacteria”), while its sister lineage, Gloeobacteria, is depauperate and contains only two closely related species of *Gloeobacter* and a metagenome-assembled genome. Here we describe a new culturable member of Gloeobacteria, *Anthocerobacter panamensis*, isolated from a tropical hornwort. *Anthocerobacter* diverged from *Gloeobacter* over 1.4 billion years ago and has low 16S identities with environmental samples. Our ultrastructural, physiological, and genomic analyses revealed that this species possesses a unique combination of traits that are exclusively shared with either Gloeobacteria or Phycobacteria. For example, similar to *Gloeobacter*, it lacks thylakoids and circadian clock genes, but the carotenoid biosynthesis pathway is typical of Phycobacteria. Furthermore, *Anthocerobacter* has one of the most reduced gene sets for photosystems and phycobilisomes among Cyanobacteria. Despite this, *Anthocerobacter* is capable of oxygenic photosynthesis under a wide range of light intensities, albeit with much less efficiency. Given its key phylogenetic position, distinct trait combination, and availability as a culture, *Anthocerobacter* opens a new window to further illuminate the dawn of oxygenic photosynthesis.

## Introduction

The rise of atmospheric oxygen is undoubtedly one of the most transformative events in Earth’s history. During the Paleoproterozoic era, oxygenic photosynthesis carried out by Cyanobacteria fueled the Great Oxygenation Event, which altered the biogeochemical cycles and fundamentally shifted the evolutionary trajectories of life on Earth^1, 2^. None of the closest living relatives of Cyanobacteria, Vampirovibrionia and Sericytochromatia^3^, are capable of oxygenic photosynthesis nor carry any intermediate photosynthetic machinery. Therefore, to understand how photosynthesis evolved over time, it is imperative to identify and examine the lineages that diverged the deepest within the Cyanobacteria tree of life.

The Phylum Cyanobacteria (defined by Garcia-Pichel et al.^4^) is composed of two extant groups: Gloeobacteria and Phycobacteria that diverged around 2 billion years ago (BYA)^5.6^. Phycobacteria encompasses >99.9% of the known cyanobacterial diversity and is sometimes also referred to as the “crown Cyanobacteria”. Gloeobacteria, on the other hand, is rather enigmatic and has only two species described thus far: *Gloeobacter violaceus* and *G. kilaueensis*. Little is known about the ecology and distribution of these two species; *G. kilaueensis* is only known from a lava cave in Hawaii^7^, and *G. violaceus* was originally isolated from a limestone rock in the Swiss alps^8^ and subsequently reported on waterfall walls in Europe and Mexico^9, 10^. The diversity of Gloeobacteria is likely much greater than these two species, as evidenced by a few distinct 16S clades from environmental samples^11^ and a recent metagenome-assembled genome (MAG), *Candidatus* Aurora vandensis, from Lake Vanda, Antarctica^12^.

Gloeobacteria is a pivotal lineage to reconstruct the early evolution of Cyanobacteria, given their many distinct, and presumably pleisiomorphic traits. In contrast to Phycobacteria, *Gloeobacter* lacks thylakoid membranes as well as a circadian clock^8, 13^. Due to the absence of thylakoids, the photosynthesis and respiratory apparatus are both located on the cytoplasmic membrane^14^. The *Gloeobacter* photosystems are also atypical^15–19^, and their phycobilisomes are bundle-shaped instead of hemidiscoidal as found in Phycobacteria^20, 21^. In addition, the carotenoid biosynthesis pathway in *Gloeobacter* is of “bacterial-type”^22, 23^, which is again different from other photosynthetic organisms. To date, a large body of research has been done on *Gloeobacter* and has significantly advanced our knowledge on Cyanobacteria biology and evolution.

However, the depauperate nature of Gloeobacteria makes it difficult to ascertain whether the idiosyncrasies of *Gloeobacter* can properly reflect the ancestral states or are instead a product of lineage-specific reductive evolution. The recent discovery of *Candidatus* Aurora MAG is an important step toward capturing the broader genomic diversity of Gloeobacteria^12^, although the fragmented nature of MAG, as well as its absence as a culturable organism, prohibit further in-depth investigations.

Here we report a new culturable Gloeobacteria species that is distantly related to *Gloeobacter*. Its complete circular genome, ultrastructure, and basic physiological properties provided new insights into the early evolution of Cyanobacteria and associated oxygenic photosynthesis.

## Results and Discussion

### Isolation of Anthocerobacter panamensis from a tropical hornwort

A new cyanobacterial culture was obtained from a surface-sterilized thallus of the hornwort (Bryophyta) *Leiosporoceros dussii* from Panama (Fig. 1). Our 16S phylogeny places this isolate (named *Anthocerobacter panamensis)* sister to a clade of uncultured environmental samples from the Arctic and Antarctic regions (e.g. Canadian tundra, Iceland, Antarctic lakes, and Patagonia) (Fig. 2A). This “polar clade” also includes *Candidatus* Aurora vandensis MAGs from Antarctica. However, *Anthocerobacter* is the only culturable strain in this clade. The 16S sequence identity between *Anthocerobacter* is around 96%, 88% and 70% when compared to polar clade members, *Gloeobacter*, and *Synechocystis* sp. PCC 6803, respectively (Fig. 2D). The equatorial origin of *Anthocerobacter* is in stark contrast to its sister polar clade, which speaks to the uniqueness of this new isolate.

**Figure 1.**
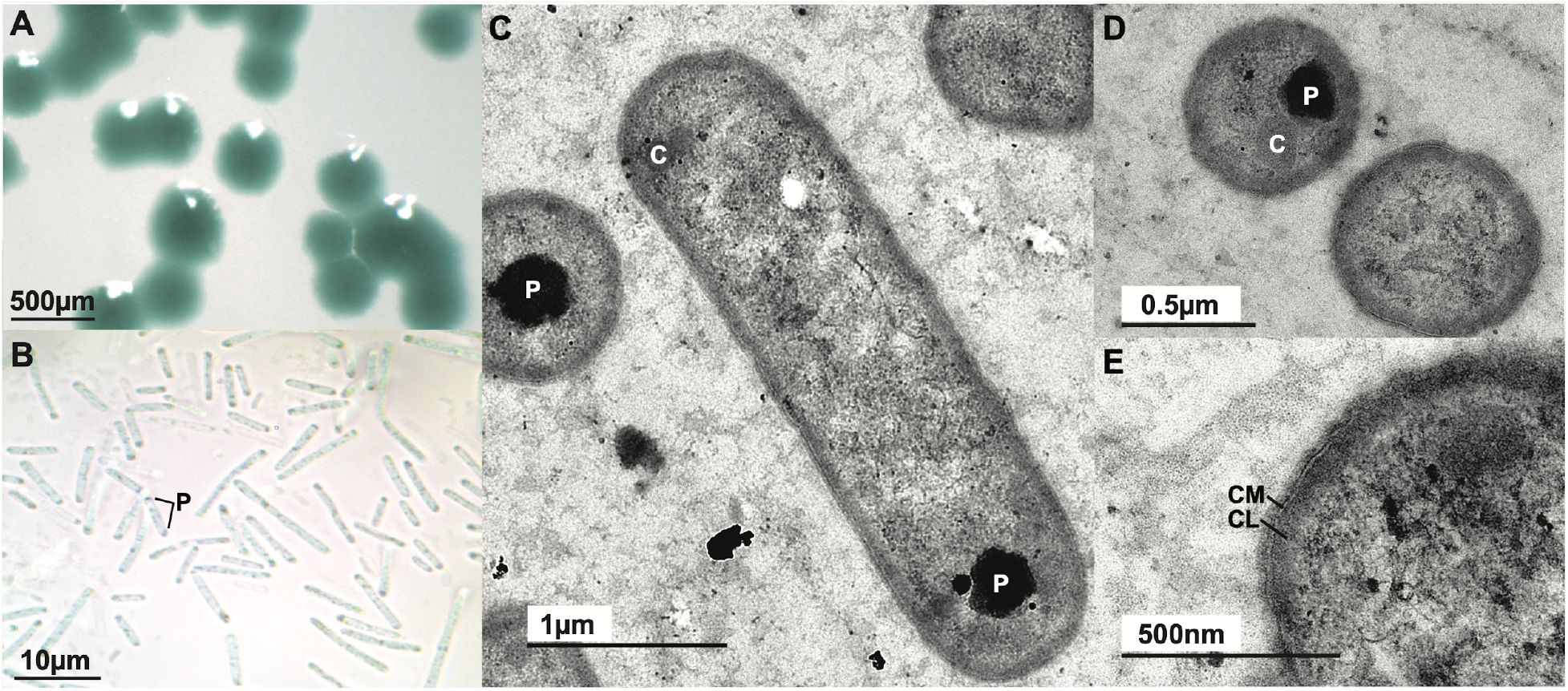
Morphology of *Anthocerobacter panamensis*. **(A)** Blue-green colonies on a solid agar plate. **(B)** Rod-shaped cells under light microscope with polyphosphate granules (P) often visible toward the cell poles. **(C-E)** TEM images of *Anthocerobacter* in longitudinal **(C)** and transverse **(D-E)** sections. No thylakoid is present. P: polyphosphate granule. C: carboxysome. CM: cytoplasmic membrane. CL: electron-dense cytoplasmic layer.

**Figure 2.**
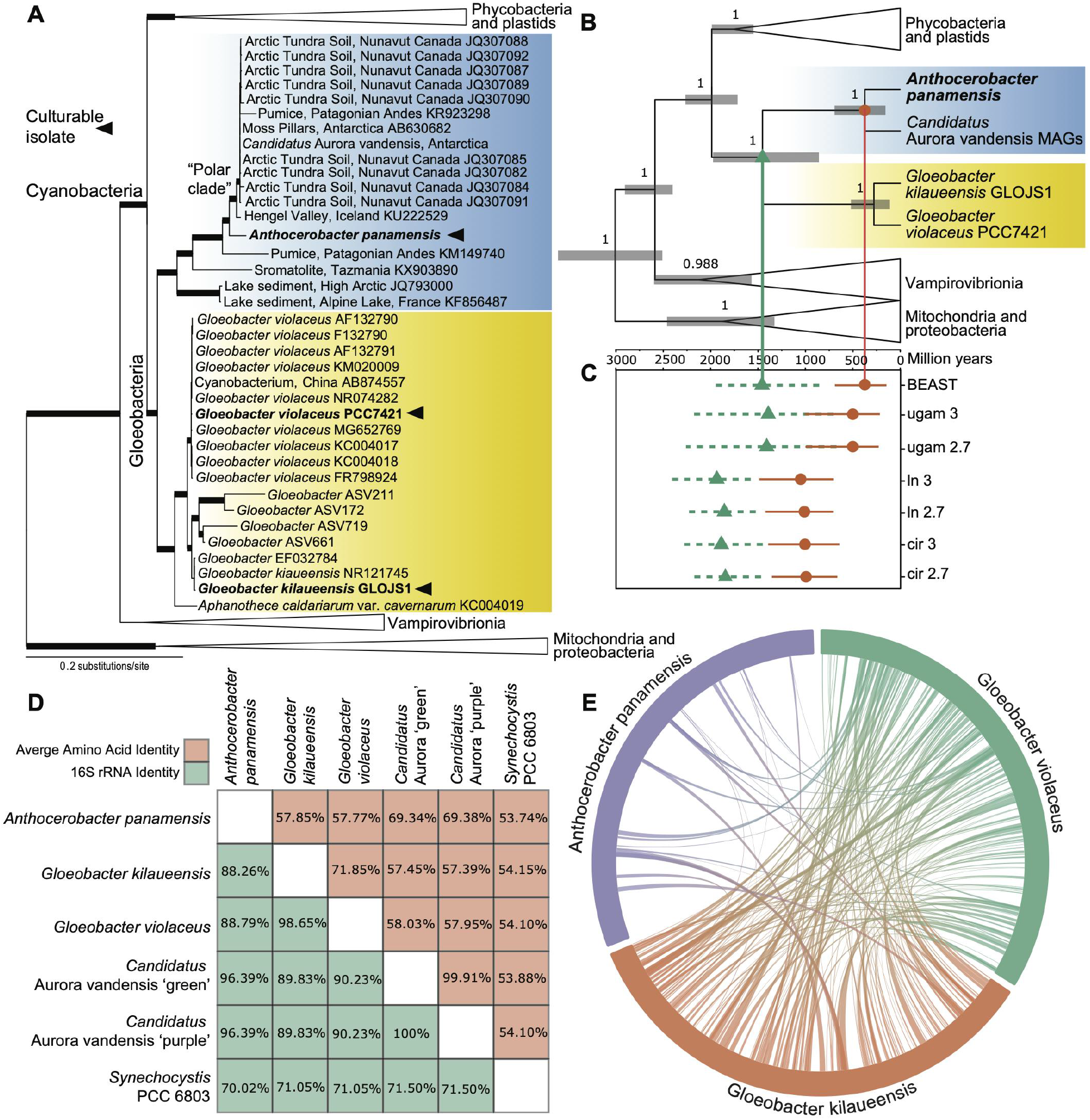
Relationship of *Anthocerobacter panamensis* and other Cyanobacteria. **(A)** *Anthocerobacter* is sister to a clade of environmental samples from the polar regions, based on 16S rRNA sequences. Thickened branches received bootstrap support >80. Arrowhead marks the strains that are available as cultures. **(B)** A time-calibrated phylogeny revealed that *Anthocerobacter* represents a deeply diverged lineage within Cyanobacteria. Values above branches are posterior probabilities. The size of collapsed wedges does not correspond to taxa number. **(C)** Comparison of divergence time estimates from **(B)** (“BEAST”) and from PhyloBayes following the calibration strategies of Baracaldo *et al.^27^*. A total of six PhyloBayes runs were carried out, using a combination of different clock models: log normal (ln), Cox–Ingersoll–Ross model (cir) and uncorrelated gamma multiplier (ugam), and two different root ages: 2.7 and 3 BYA. The orange and green lines are the 95% HPD of age estimates for *Anthocerobacter-Candidatus* Aurora and *Anthocerobacter-Gloeobacter* splits, respectively. **(D)** *Anthocerobacter* has low pairwise 16S rRNA identities and low AAI with other cyanobacterial species. *Candidatus* Aurora vandensis “green” and “purple” are MAGs from separate sediment layers. **(E)** Very little genomic synteny exists between *Anthocerobacter* and *Gloeobacter spp*.

Because all hornwort species have symbiotic associations with filamentous nitrogen-fixing Cyanobacteria^24, 25^, it raises the question of whether *Anthocerobacter* could be a major symbiont. We suggest this is unlikely to be the case as *Anthocerobacter* is not filamentous (Fig. 1B), and its genome lacks nitrogenase genes while nitrogen fixation is the selective advantage in plant-Cyanobacteria symbiosis. Recently, the microbiome of *L. dussii* was profiled using 16S amplicon sequencing^26^, but no trace of *Anthocerobacter* could be found (though interestingly, a unique clade of *Gloeobacter* was uncovered therein: ASV172, ASV719, and ASV661 in Fig. 2A). We therefore speculate that *Anthocerobacter* could be a soil bacterium that happened to survive our surface sterilization, an endophyte that inhabits hornwort intercellular space, or a freeloader inside symbiotic cavities with other functional nitrogen-fixing cyanobionts. Future studies are needed to clarify the ecology and natural history of *Anthocerobacter*.

### Anthocerobacter represents a novel and deeply branched cyanobacterial lineage

To corroborate the phylogenetic placement of *Anthocerobacter* and to estimate the timing of divergence, we compiled a 12-gene matrix based on the dataset of Shih *et al.*^6^. These genes are slow-evolving, and are shared between plastid and mitochondria genomes, thereby allowing for cross-validations to reuse and link the same calibration across multiple nodes. As in the 16S phylogeny, we recovered the same branching order among *Anthocerobacter, Candidatus* Aurora, *Gloeobacter*, and Phycobacteria with strong support (Fig. 2B). It should be noted that our topology is different from the 16S tree of Grettenberger *et al.*^12^, which resolved *Candidatus* Aurora sister to *Gloeobacter* + Phycobacteria, but is consistent with their 37-gene phylogeny. Given the high branch support from both of our datasets, we believe our result, (((*Anthocerobacter,Candidatus* Aurora),*Gloeobacter*), Phycobacteria), represents the most plausible relationship.

The chronogram from BEAST suggested that *Anthocerobacter* diverged from *Candidatus* Aurora at around 374.74 million years ago (MYA) (95% highest posterior density (HPD): 155.86–694.79 MYA), and from *Gloeobacter* around 1.45 BYA (95% HPD: 857.64–1,975.35 MYA)(Fig. 2B). Because molecular dating analyses can be strongly influenced by calibrations, we further explored the robustness of our time estimates by adopting a suite of different calibration schemes, models, and data matrices made by Baracaldo *et al.*^27^. Comparable but generally deeper divergences were obtained, and the results are summarized in Fig. 2C. Taken together, our phylogenetic and molecular clock analyses indicate that *Anthocerobacter* is currently the sole culturable representative of a novel cyanobacterial lineage, which split from *Gloeobacter* over 1.4 billion years ago.

### Anthocerobacter genome

Nanopore sequencing resulted in one circular genome assembly of 4,187,797bp and a GC content of 55.51%. The Average Amino-acid Identity (AAI) is around 57% when compared to *Gloeobacter* and 69% to *Candidatus* Aurora (Fig. 2D). A total of 4,166 protein-coding genes were predicted, 1,805 of which can be functionally annotated. Protein family circumscription based on 103 cyanobacterial species identified a total of 18,419 orthogroups. Of these, 2,818 gene families are present in *Anthocerobacter*, 35 are species-specific, and 140 are unique to *Anthocerobacter* + *Candidatus* Aurora (Table S1; Fig. S1). A total of 454 predicted genes in *Anthocerobacter* were singletons that could not be assigned to any orthogroup. Furthermore, we found very little synteny between *Anthocerobacter* and *Gloeobacter spp.*, consistent with the deep divergence between the two genera (Fig. 2E).

### Anthocerobacter lacks thylakoids

One of the most distinctive features of *Gloeobacter* is its lack of thylakoids^8, 9^. While the ultrastructure of *Candidatus* Aurora is not available, analysis of its metagenome found a lack of the *VIPP1* (vesicle inducing protein in plastid 1) gene^12^, which is essential for biogenesis of thylakoid membrane^28^. The availability of *Anthocerobacter* as a pure culture allowed us to directly examine it using transmission electron microscopy (TEM). We found no trace of thylakoids (Fig. 1C-E), nor a *VIPP1* homolog in the genome. This result reaffirmed that the lack of thylakoid is likely a general feature of Gloeobacteria, and thylakoids were an innovation of Phycobacteria.

### Spectroscopy reveals unique features of Anthocerobacter

In a stark contrast to the purple to brown color of *Gloeobacter*^7^, ^9^, *Anthocerobacter* are blue-green and has no apparent red hue (Fig. 1A). To better understand the pigment composition, we characterized and compared *Anthocerobacter*’s absorption and fluorescence emission spectra with those of *Gloeobacter violaceus*, and *Synechocystis* sp. PCC 6803 (a model unicellular species from Phycobacteria; hereafter Syn6803). In the whole cell absorption spectra (Fig. 3A), *Anthocerobacter* lacks clear peaks representing phycoerythrin (PE), which is different from *G. violaceus*^29^ but consistent with the absence of PE genes in the *Anthocerobacter* genome (see below). Compared to Syn6803, which shows predominant Soret and Qy bands of chlorophyll (Chl) *a* (440 nm and 682 nm), the absorption at 682 nm in *Anthocerobacter* is lower, not showing a pronounced peak (Fig. 3A). The peak at 635 nm from phycocyanin (PC) is higher in *Anthocerobacter* than in Syn6803 at 629 nm, though it is common that the absorption peaks of the same phycobiliproteins differ from species to species^21, 30, 31^. The absorption feature at 495 nm is from carotenoids. Because *Anthocerobacter* has no PE nor phycoerythrocyanin (PEC) identified in the genome, the 582 nm feature, which is absent in Syn6803, is from an unknown source, possibly from carotenoid-binding proteins. In *G. violaceus*, this wavelength region is overlaid with the absorption of PE in the whole cell absorption spectra and was therefore not discussed previously^29^.

**Figure 3.**
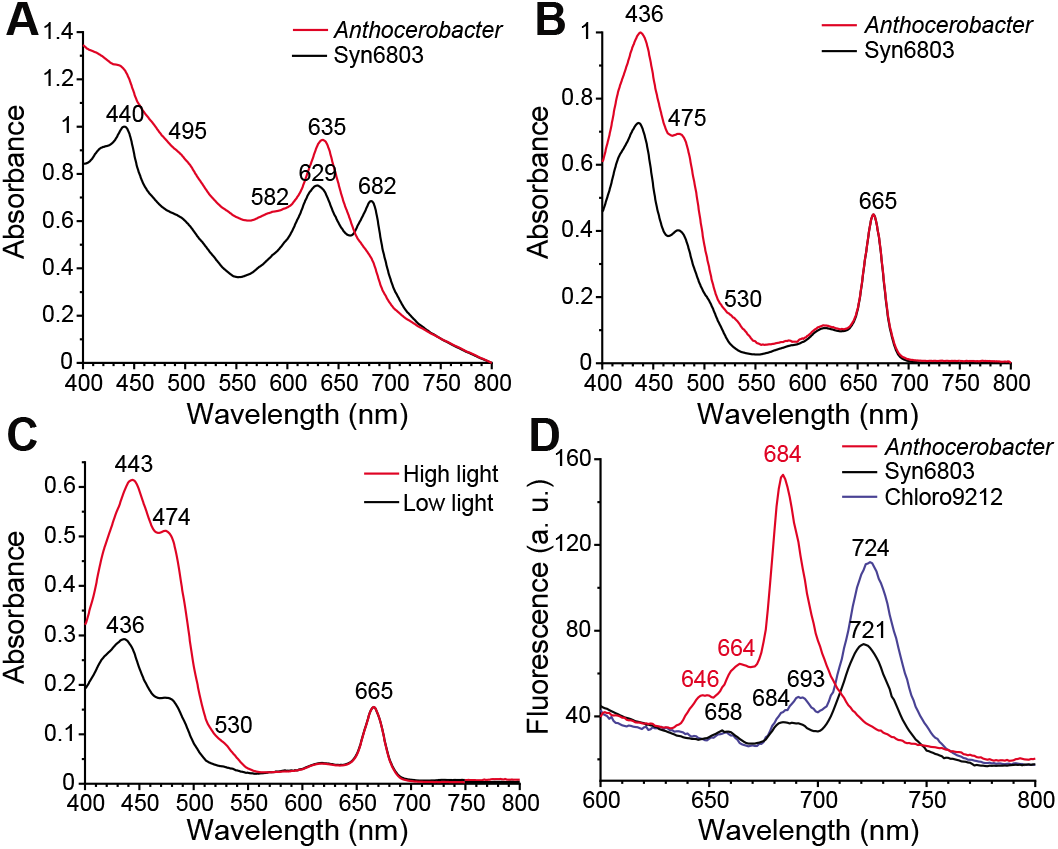
Spectral analyses of *Anthocerobacter* in comparison with other Cyanobacteria. Absorption spectra of **(A)** whole cells, **(B)** pigment extraction in methanol, and **(C)** pigment extraction in methanol of *Anthocerobacter* in high and low light (100 and 10 μmol photons m^−2^s^−1^) **(D)** Fluorescence spectroscopy was taken at 77K with excitation wavelength at 440 nm using whole cells of *Anthocerobacter*, Syn6803, and *Chlorogloeopsis fritschii* PCC 9212 (Chloro9212). The absorbance spectra were normalized at 750 nm in (A) and at 665 nm in (B) and (C). Specific wavelengths were labeled above the peaks.

The absorption spectra from methanol-extracted pigments (Fig. 3B) shows that *Anthocerobacter* contain only Chl *a* (665 nm and 436 nm) and no other chlorophylls, and the Qy band (665 nm) matches perfectly with the Qy band from Chl *a* in Syn6803. However, although both strains show features of carotenoids at 475 nm, *Anthocerobacter* have stronger absorption at 436 nm and an additional absorption feature around 530 nm, indicating it has one or more carotenoids with long-wavelength absorption. A long-wavelength absorbing carotenoid was also identified in *G. violaceus*, and its absorption maximum is similar to oscilloxanthin^14, 29^. At high-light condition (100 μmol photons m^−2^s^−1^), *Anthocerobacter* produced more carotenoids (absorption peaks at wavelengths 443 nm, 474 nm, and 530 nm) per unit of Chl *a* (Fig. 3C and 5B). Further analyses are required to characterize the species and quantities of carotenoids in *Anthocerobacter*.

Finally, 77K fluorescence spectroscopy was used to measure fluorescence emission from chromophore-bound protein complexes such as phycobilisomes, photosystem I (PSI) and II (PSII). In *Anthocerobacter*, phycobiliproteins are the predominant light-harvesting complexes, resulting fluorescence emission largely coming from phycobiliproteins even though the excitation wavelength (at 440 nm) is targeted for chlorophylls. Three emission peaks are observed in *Anthocerobacter* and assigned as PC (646 nm), AP (664 nm), and PSII (684 nm) (Fig. 3D). The peak at 684 nm is asymmetric, indicating that the second peak of PSII is likely present but not strong enough forming a separate peak, a pattern also found in *G. violaceus*^15^, ^29^. The long-wavelength emission from PSI is absent in *Anthocerobacter*, which is different from Syn6803 (and other Phycobacteria), but similar to *G. violaceus* (Figure 3D)^15, 29^. Previous studies in *G. violaceus* concluded that the absence of long-wavelength emission beyond 700 nm is due to the lack of long-wavelength Chl *a* in PSI^29^, which is possibly the situation in *Anthocerobacter* as well. These measurements indicate that the spectral properties of *Anthocerobacter* are distinct from Syn6803 but largely similar to *G. violaceus* with the exception that *Anthocerobacter* lacks PE.

### Anthocerobacter lacks cyanobacterial circadian clock genes

Circadian clocks enable organisms to adjust their physiological activities according to extrinsic daily changes such as light, temperature, and humidity. In Cyanobacteria, environmental signals are transmitted through redox sensitive proteins to core oscillators (KaiA, KaiB and KaiC), followed by output pathways to regulate physiological activities^32^. We found that the *Anthocerobacter* genome is missing the core oscillator KaiABC genes, similar to what was reported in *Gloeobacter*^13^. This implies that the cyanobacterial circadian clock most likely evolved in the common ancestor of Phycobacteria.

### The carotenoid biosynthesis pathways of Anthocerobacter and Gloeobacter differ

Carotenoids are an integral part of light harvesting complexes and play key roles in photoprotection in conjunction with orange carotenoid proteins (OCP). While *Anthocerobacter* OCP is in the OCPX clade (defined by Bao *et al*.^33^) like *Gloeobacter* (Fig. S2, Table S2), the pathway through which carotenoids are made differs between the two lineages. The first step of carotenoid biosynthesis is the condensation of two geranylgeranyl pyrophosphate into phytoene by phytoene synthase (CrtB)(Fig. S3). Following this, phytoene is converted to lycopene through two different routes depending on the organisms. The “plant-type” pathway, which can be found in all Cyanobacteria (except *Gloeobacter)* and plants, relies on three separate enzymes: phytoene desaturase (CrtP), ζ-carotene desaturase (CrtQ), and cis-carotene isomerase (CrtH)^34^. By contrast, the “bacteria-type”, found in *Gloeobacter*, anoxygenic photosynthetic bacteria, and fungi, uses only a single phytoene desaturase (CrtI)^22, 23^. Interestingly, we found that both *Anthocerobacter* and *Candidatus* Aurora have the genetic chassis for the plant-type carotenoid biosynthesis, instead of the bacteria-type reported in *Gloeobacter* (Fig. S3). This implies that the origin of the plant-type carotenoid biosynthesis pathway should be pushed back to the ancestor of all Cyanobacteria, with *Gloeobacter* later substituting it with the bacterial pathway. While we cannot entirely rule out that *Anthocerobacter* horizontally acquired the plant-type pathway from Phycobacteria, the phylogenetic positions of *Anthocerobacter crtP, crtQ*, and *crtH* are consistent with the species phylogeny (Fig. S4 and S5).

### Anthocerobacter has a unique phycobilisome composition

Cyanobacteria use phycobilisomes as the light harvesting antennae to capture and transfer light energy to photosystems. Typically, phycobilisomes are fan-shaped and composed of the allophycocyanin core (AP) and several peripheral proteins: phycocyanin (PC) and phycoerythrin (PE) or phycoerythrocyanin (PEC)^21, 30^. One notable exception is *Gloeobacter*, whose phycobilisomes are bundle-shaped and consist of AP, PC, and PE^20, 21^. Compared to *Gloeobacter, Anthocerobacter* further lacks homologs for PE (and PEC), a situation also reported from *Candidatus* Aurora MAG (Fig. 4)^12^. On the other hand, most of the AP and PC subunits are present in *Anthocerobacter*, which is consistent with our absorption and fluorescence emission spectral data (Fig. 3A and D). The missing AP and PC subunits are ApcD, ApcF, and CpcD, although the mutations of which did not appear to affect the phycobilisome function in *Synechococcus* sp. PCC 7002^35, 36^ or in Syn6803^37, 38^.

**Figure 4.**
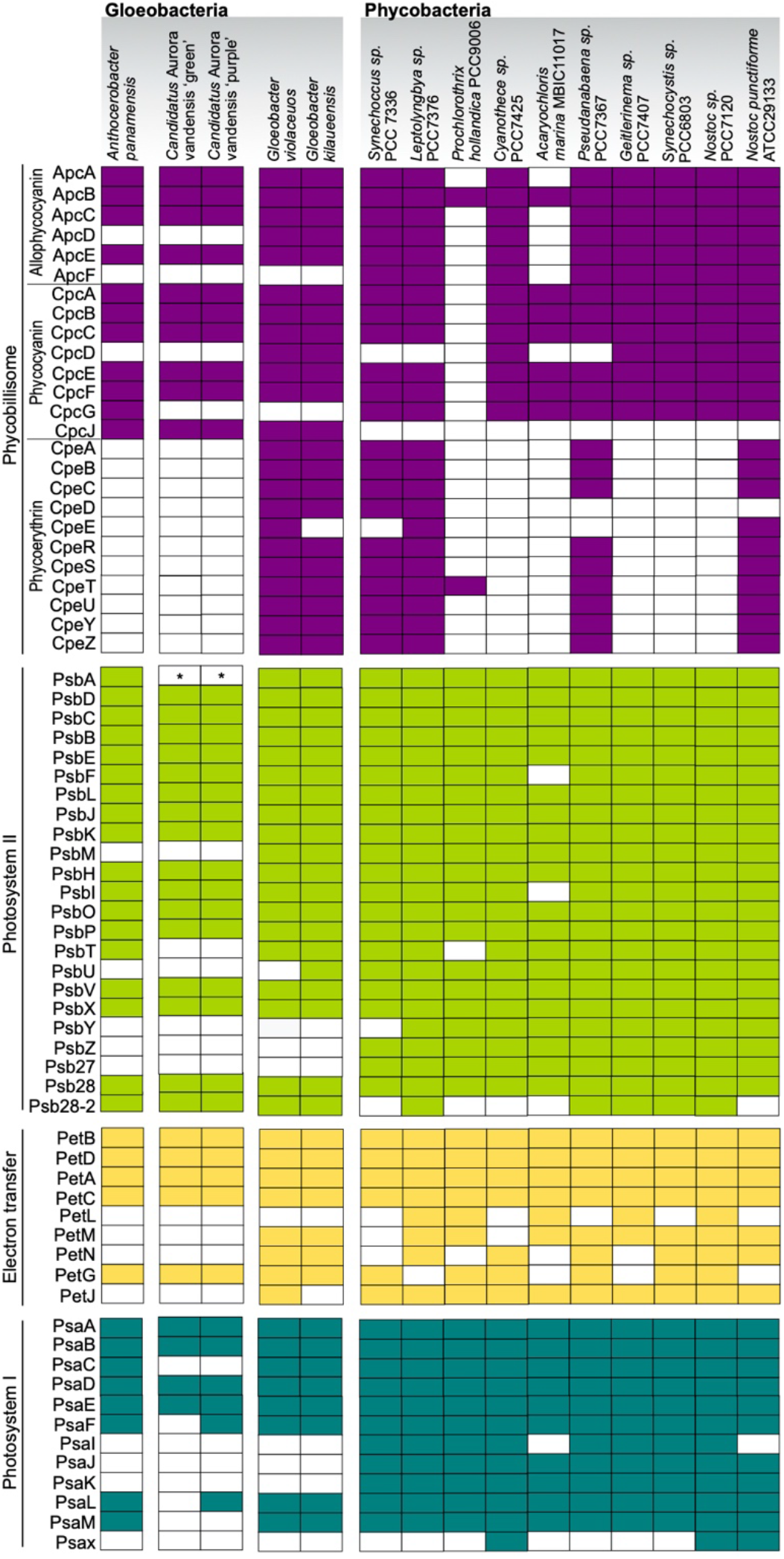
Comparison of phycobilisome and photosystem subunits among *Anthocerobacter, Candidatus* Aurora, *Gloeobacter*, and other Cyanobacteria. Filled and empty boxes indicate gene presence and absence respectively. *Candidatus* Aurora “green” and “purple” are MAG from separate sediment layers. *\*psbA* was not found in Aurora MAG but on separate contigs that could not be properly binned.

**Figure 5.**
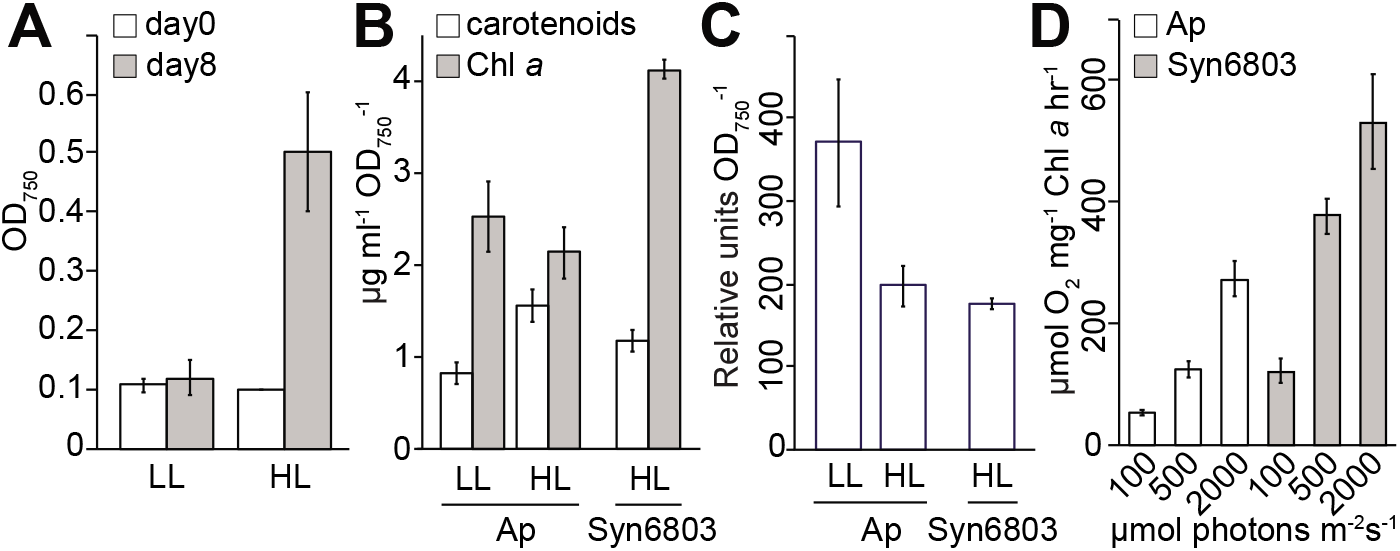
Measurements of growth, pigment analyses, and oxygen evolution. *Anthocerobacter* was grown in high light (HL, 100 μmol photon m^−2^s^−1^) and low light (LL, 10 μmol photon m^−2^s^−1^) conditions for 8 days. **(A)** The OD_750_ values at the beginning of inoculation (Day 0) and the end of cultivation (Day 8) were recorded for estimation of doubling time. Quantifications of **(B)** Chl *a* and carotenoids, and **(C)** phycobiliproteins were performed using *Anthocerobacter* (Ap) cells grown in LL and HL on Day 8. *Synechocystis* PCC6803 (Syn6803) grown under HL was used as a comparison. All the calculations were based on three biological replicates. **(D)** Rates of oxygen evolution (Y-axis) over different light intensities (μmol photons m^−2^s^−1^). Three biological replicates were used for all the measurements.

Another peculiar feature of *Anthocerobacter* phycobilisomes is the presence of *cpcG*, which encodes for a rod-core linker protein. This protein is critical in phycobilisome assembly and energy transfer from PC to AP but is absent in *Gloeobacter* and *Candidatus* Aurora. In *Gloeobacter*, a different linker protein, CpcJ, was used instead of CpcG^39^. Interestingly, this *Gloeobacter*-specific *cpcJ* gene is also present in *Anthocerobacter*, suggesting that *Anthocerobacter* has two different types of rod-core linker (CpcG and CpcJ) for connecting PC and AP (Table S2). In other words, *Anthocerobacter* may have a considerably different phycobilisome from that of *Gloeobacter*, which is already distinct among Cyanobacteria. Furthermore, our phylogenetic analysis of phycobilisome linker proteins placed *Anthocerobacter cpcG* sister to the rest of cyanobacterial homologs (Fig. S6), suggesting that the ancestor of Cyanobacteria very likely had the CpcG linker.

### Anthocerobacter has a reduced set of photosystem subunits

With respect to photosystems, *Anthocerobacter* is similar to *Gloeobacter* in that they both lack several highly conserved genes encoding PsbY, PsbZ, and Psb27 at photosystem II (PSII), and PsaI, PsaJ, PsaK, PsaX at photosystem I (PSI)(Fig. 4). These subunits are involved in electron transfer (e.g. PsbZ)^40^, or the formation and stabilization of photosystems (e.g. Psb27, PsaI, and PsaJ)^41–45^. *Anthocerobacter* also lacks the known pathway genes for synthesizing phylloquinone/menaquinone (KEGG M00116), which is a secondary electron acceptor of PSI; such absence is also shared with *Gloeobacter^18^*.

Interestingly, we found even more genes encoding photosystem subunits are missing in *Anthocerobacter* (and to some extent *Candidatus* Aurora as well) compared to *Gloeobacter*, including PsbM, PsbU, PetM, and PetN. PsbM is localized in the PSII center and plays a key role in PSII dimer formation and stability^46^. PsbU is needed for making a thermally stable oxygen-evolving complex, and aids electron transport and energy transfer from phycobilisome to PSII^47, 48^. At the electron transport chain, PetN is necessary for the proper function and assembly of the cytochrome b6f complex at least in Syn6803 and tobacco^49^. While PetM mutations in Syn6803 did not impair the cytochrome b6f function, the amounts of phycobilisomes and PSI decreased^50^. In all, *Anthocerobacter* has one of the most reduced sets of photosynthesis subunits among Cyanobacteria, which prompted us to further investigate the physiological properties of *Anthocerobacter* and compare them with existing cyanobacterial strains.

### Anthocerobacter is able to grow photoautotrophically and evolve oxygen, but slowly

The growth of *Anthocerobacter* was measured under low light (LL) and high light (HL) conditions (10 and 100 μmol photons m^−2^s^−1^, respectively) in the medium without supplement of carbon sources. *Anthocerobacter* can grow photoautotrophically under both light intensities, with a faster rate under HL (Fig. 5A). The pigment analyses show that under HL, the Chl *a*:phycobiliprotein ratio is about half in *Anthocerobacter* as in Syn6803 (Fig. 5B-C). *Anthocerobacter* produces less Chl *a* and phycobiliproteins but more carotenoids in HL than in LL, probably because photoprotection is turned on. A similar change from lower to higher light intensity was also observed in *Gloeobacter*^15^.

Compared to other model Cyanobacteria, *Anthocerobacter* is very slow growing; its doubling time is estimated around 67 h under HL and over 8 days under LL (Fig. 5A). By comparison, Syn6803 doubles in around 8 hours under the same HL condition. The reported doubling time for *Gloeobacter* ranges from 73 h to 17 days^14, 51^, depending on the culturing conditions. The slow growth of *Anthocerobacter* likely results from its inefficient photosynthetic apparatus and/or the inability to tolerate high light intensities, which could be caused by the fact that they lack accessory subunits of PSI and PSII (Fig. 4).

Finally, we demonstrated that *Anthocerobacter* does produce oxygen, and the level of which increased with light intensity (from 100, 500, to 2000 μmol photons m^−2^s^−1^) (Fig. 5D). In general, *Anthocerobacter* has about half of the oxygen evolution rates compared to Syn6803 under the same conditions. While *Gloeobacter* was not included in our experiments, we could cross-reference the results with those of Koyama *et al.^52^*, which reported a similar difference in oxygen evolution rates between Syn6803 and *G. violaceus*. In other words, despite having even fewer photosynthesis subunits than *Gloeobacter, Anthocerobacter* can nevertheless carry out comparable oxygenic photosynthesis. These results in *Anthocerobacter* may contribute to determination of the minimal and essential photosynthetic subunits required for photoautotrophic growth in nature.

### *Comparison with* Candidatus *Aurora*

Very recently, *Candidatus* Aurora was described from a metagenome assembly^11^, which based on our phylogenetic analysis belongs to the same lineage as *Anthocerobacter* (Fig. 2). There are however several key differences between the two. First, the assembly of *Candidatus* Aurora MAG is highly fragmented and incomplete, making it difficult to confidently conclude gene presence/absence and discuss the broader implications on cyanobacterial evolution. In fact, *psbA* (which encodes PSII reaction center D1 protein) was not present in Aurora MAG, but instead found on a separate small contig that was not binned properly. Here, with a complete circular genome of *Anthocerobacter*, we are able to fully describe the gene content. A case in point is the phycobilisome linker protein CpcG. Because it is missing in *Gloeobacter* genomes as well as Aurora MAG, *cpcG* gene would be inferred to originate in Phycobacteria. However, we found a clear *cpcG* homolog in *Anthocerobacter* that is phylogenetically sister to the rest of cyanobacterial *cpcG* (Fig. S6). This suggests that *cpcG* is probably ancestral to all Cyanobacteria (but subsequently lost in *Gloeobacter)* and raises the possibility that its absence in Aurora MAG might be due to incomplete assembly.

Second, the existence of *Candidatus* Aurora had been hinted by earlier environmental samples from Arctic and Antarctic regions, which share highly similar 16S sequences (>99% identity; Fig. 2)^11^. *Anthocerobacter*, on the hand, was not known from any environmental sample that we are aware of. In addition, as an isolate from a tropical hornwort, *Anthocerobacter* shows a contrasting distribution pattern from its sister polar clade (Fig. 2).

Most importantly, *Candidatus* Aurora is not an isolated culture, rendering further studies difficult if not impossible. The fact that *Anthocerobacter* is the only culturable organism in this lineage allowed us to characterize the morphology and physiology of this representing strain in detail. The lack of thylakoid in *Anthocerobacter* was observed by TEM (Fig. 1), which is much more convincing than basing on the absence of *VIPP1* homolog alone in *Candidatus* Aurora^11^, especially considering gene absence could also be due to incomplete MAG assembly. Our spectral and growth experiments also provided important insights into the biology of *Anthocerobacter* that are otherwise impossible to obtain for *Candidatus* Aurora. For example, *Anthocerobacter* lacks long-wavelength emission from PSI similar to *Gloeobacter*, suggesting that this feature is not an idiosyncrasy of *Gloeobacter* but deeply shared across Gloeobacteria. In addition, *Anthocerobacter* was experimentally demonstrated capable of oxygen evolution and photoautotrophic growth, which are not possible to be verified in *Candidatus* Aurora. While there is no doubt that the discovery of *Candidatus* Aurora is significant, *Anthocerobacter*, owing to its culturable nature, is the one that can advance our understandings of this key cyanobacterial lineage.

### Conclusion

*Anthocerobacter* and *Candidatus* Aurora together represent a novel and ancient lineage of Cyanobacteria, which diverged from *Gloeobacter* over 1.4 billion years ago. The deep divergence and availability as a culture enable us to use *Anthocerobacter* to revisit the early evolution of Cyanobacteria. One key question we were able to address is whether the many unique features in *Gloeobacter* are indeed characteristic of the entire Gloeobacteria lineage. On one hand, we found that similar to *Gloeobacter, Anthocerobacter* lacks thylakoids, circadian clock oscillators, nitrogenase, canonical pathway for phylloquinone synthesis, long-wavelength PSI fluorescence, as well as many conserved photosynthesis subunits (summarized in Fig. S7). The most parsimonious interpretation is that the origins of these components took place in the most recent common ancestor (MRCA) of Phycobacteria, and the absence of which likely define Gloeobacteria. Conversely, the carotenoid biosynthesis pathway in *Anthocerobacter* is not the bacterial-type as found in *Gloeobacter*, but typical of the rest Cyanobacteria, implying that the plant-type pathway likely evolved in the MRCA of Cyanobacteria and not of Phycobacteria (Fig. S7). *Anthocerobacter* also possesses several unique features, such as its phycobilisome composition and an even further reduced collection of photosystem subunits. Nevertheless, we were able to demonstrate that *Anthocerobacter* can indeed grow photoautotrophically and evolve oxygen across a range of light intensities, although less efficiently compared to other cyanobacterial strains. In summary, *Anthocerobacter* is an important addition to the depauperate Gloeobacteria and will facilitate future investigations into the origin of Cyanobacteria and oxygenic photosynthesis.

### Taxonomic treatment

#### Anthocerobacter

F.-W. Li gen. nov. (Fig. 1)

Differ from genera of Chroococcales, Chroococcidiopsidales, Nostocales, Oscillatoriales, Pleurocapsales, Spirulinales, and Synechococcales in lacking thylakoid membranes. Most similar to *Gloeobacter* in Gloeobacterales but lacks phycoerythrin.

#### Etymology

An.tho.cero.bact’er. N.L. neut. n. Anthocerotophyta, hornworts; N.L. masc. n. bacter, a small rod; N.L. masc. n. Anthocerobacter, a rod from hornworts.

#### Type species

*Anthocerobacter panamensis* F.-W. Li sp. nov.

#### Anthocerobacter panamensis

F.-W. Li sp. nov. (Fig. 1)

The cells are rod-shaped, 0.5-1 μm wide and 2-10 μm long. Thylakoid membranes are absent. Carboxysomes are present, and polyphosphate granules often appear toward the cell poles. The major pigments are allophycocyanin, phycocyanin, and chlorophyll *a*. No phycoerythrin nor phycoerythrocyanin is present. Colonies are generally light to dark green in both liquid and solid BG11 medium, and never purple, orange, or brown like in *Gloeobacter*. Growth can occur under 10 - 100 μmol photons m^−2^s^−1^ light intensities. No nitrogenase genes are present and cells do not grow without supplementing combined nitrogen. The type strain (C109) was originally isolated from a surface-sterilized hornwort *Leiosporoceros dussii* from Coclé, Panama (8°37’34.6” N, 80°08’13.7” W). GC content of the type strain is 55.51%.

#### Etymology

pa.na.men’sis. N.L. masc./fem. adj. panamensis, of or pertaining to Panama where the type strain was isolated.

#### Holotype

The type strain C109 was deposited at Bailey Hortorium (BH) under the accession xxx, and American Type Culture Collection (no. xxx).

## Materials and Methods

### Culturing

The hornwort *Leiosporoceros dussii* (Steph.) Hässel was collected in Coclé, Panama (8°37’34.6” N, 80°08’13.7” W). Sterilization of the plant thalli and initial culturing of the isolate (type strain C109) followed the protocol of Nelson *et al.^53^. Synechocystis* sp. PCC 6803 was a gift from Dr. Hsiu-An Chu at Institute of Plant and Microbial Biology, Academia Sinica, Taiwan, and *Chlorogloeopsis fritschii* PCC 9212 was obtained from the Pasteur Culture Collection^54^. *Anthocerobacter, Chlorogloeopsis fritschii* PCC 9212, and *Synechocystis* sp. PCC 6803 were grown in the B-HEPES growth medium^55^, a modified BG11 medium containing 1.1 g L^−1^ HEPES (final concentration) with the pH adjusted to 8.0 with 2.0 M KOH. Cool white LED light provided continuous illumination at either 10 or 100 μmol photons m^−2^s^−1^ in a 30°C growth chamber supplemented with 1% (v/v) CO2 in air. The doubling time is estimated based on the change of OD750 absorption over time.

### DNA extraction

Cyanobacteria colonies from either liquid cultures or solid BG11 plates were centrifuged at 17,000 x g for 2 mins to pellet the cells. The cells were ground with a pestle in liquid nitrogen and resuspended in a prewarmed 2X CTAB solution with 1% beta-mercaptoethanol. The mixture was incubated at 65°C for 1h, with gentle mixing every 15 min. Equal volume of 24:1 chloroform:isoamyl alcohol was added to the samples twice, each time with 5 min mixing on a Labnet Mini Labroller and 5 min centrifugation at 17,000 x g. The supernatant was transferred to a new tube by wide bore pipette tips to maintain DNA integrity. DNA was precipitated with an equal volume of isopropanol and pelleted for 30 min at 17,000 x g and 4°C. The DNA pellets were washed twice with 70% ethanol, then air dried in a sterile air hood and resuspended in Tris–HCl pH 7.5, before treated with RNase for 1 h at 37°C.

### Nanopore and Illumina sequencing

Genomic DNA was sequenced on both Oxford Nanopore MinION and Illumina platforms. Nanopore libraries were prepared using the Ligation Sequencing kit (SQK-LSK109) and sequenced on two MinION R9 flowcells (FLO-MIN106D) for 60 hours. In one MinION run, the focal sample was multiplexed with another cyanobacteria sample and barcoded by Native Barcoding Expansion Kit 1-12, while the other run contains only the focal sample. Basecalling was carried out using Guppy v3.1.5 with the high accuracy flip-flop mode. The Illumina library was prepared and sequenced on NextSeq500 by MiGS (Microbial Genome Sequencing Center). The raw nanopore and Illumina reads are deposited at NCBI SRA under the accessions SRR12713672 and SRR12713672.

### Genome assembly and annotation

The basecalled nanopore reads were first filtered to remove reads shorter than 10kb, and then *de novo* assembled using Flye v2.4.1^56^ with the --plasmids flag on, followed by polishing by medaka (https://github.com/nanoporetech/medaka). The resulting assembly contained three circular contigs. Based on the PATRIC taxonomic assignments^57^, one has the closest affinity to *Gloeobacter*, while the other two belong to *Pseudomonas fluorescens*, indicating contaminations were introduced during liquid culture (which was later confirmed by light microscopy). We therefore started a new culture for generating Illumina data. After trimming by TRIMMOMATIC (min score=25, min length=36)^58^, the resulting Illumina reads had 91.74% mapping rate against the cyanobacterial contig (based on the bwa mem aligner^59^), suggesting most (if not all) contaminations were removed. Genome polishing with Illumina reads was done by pilon^60^ for 4 iterations. Structural and functional annotation was performed using PATRIC^57^, with additional KEGG annotation by BlastKOALA^61^. The genome assembly is deposited at NCBI under BioProject PRJNA665722.

### Genome comparisons

Pairwise 16S sequence divergence and Average Amino-acid Identity (AAI) were calculated by seqinr v3.6-1^62^ and CompareM v0.1.1 (https://github.com/dparks1134/CompareM), respectively. We used Orthofinder v2.3.12^63^ to classify families of orthologous genes in the genomes of *Anthocerobacter* and 102 other cyanobacterial species. UpSetR^64^ was used to summarize the shared and unique orthogroups found in *Anthocerobacter*. To infer synteny between *Anthocerobacter* and the two *Gloeobacter* species, MCScanX^65^ was used with Diamond^66^ as the search engine. The syntenic relationship was then visualized in AccuSyn (https://accusyn.usask.ca).

### Phylogenetic analyses and divergence time estimates

We first compiled a 16S data matrix that includes major Cyanobacteria lineages, outgroups^6^, and available environmental sequences with high sequence similarities to *Gloeobacter* and *Anthocerobacter*^11^, ^12^. Alignment was done by PASTA v3^67^, and maximum-likelihood phylogeny was inferred using IQTREE v1.6.11 with 1,000 ultrafast bootstrap replicates^68^.

Next we integrated our data with previous molecular clock analyses on Cyanobacteria: Shih *et al.^6^* and Baracaldo *et al.^27^*. The dataset from Shih *et al.^6^* is composed of 11 concatenated protein sequences *(atpA, atpB, atpE, atpF, atpH, atpI, rpl2, rpl16, rps3, rps12*, and elongation factor Tu) and 16S nucleotide sequences. We used the same age calibrations and substitution models (CpREV for proteins and GTR+Gamma for 16S) as in Shih *et al.^6^*. Phylogeny and divergence time were inferred using BEAST v.1.8.3^69^ on CIPRES Science Gateway (v3.3)^70^. Five MCMC were run in total, and for each chain samples were taken every 10,000 generations. The posterior distributions were inspected in Tracer v.1.7.1 to ensure proper mixing and convergence. The resulting trees from five runs were combined by LogCombiner and summarized by TreeAnnotator^69^.

The dataset from Baracaldo *et al.^27^* is composed of 8 protein-coding genes (*atpA*, *atpB, petB, psaC, psbA, psbD, rbcL*, and S12). Following the protocol of Baracaldo *et al.^27^*, PhyloBayes^71^ was used to estimate divergence time, with CAT-GTR+Gamma substitution model, three different relaxed molecular clock models: log normal (ln), Cox–Ingersoll–Ross model (cir) and uncorrelated gamma multiplier (ugam), and two different root priors: (1) 95% of the prior distribution falls between 2.32 and 2.7 BYA and (2) between 2.3 and 3 BYA. The sequence alignments, input files for BEAST and PhyloBayes, as well as the resulting tree files can be found at https://gitlab.com/NasimR/cyanobacteria.

Phylogenetic inferences were also done for selected orthogroups using IQTREE v1.6.11 with 1,000 ultrafast bootstrap replicates and automatic model selection^68^.

### Phylogeny of orange carotenoid proteins (OCP)

OCP is composed of two domains, N-terminal domain (NTD) and C-terminal domain (CTD), connected by a flexible loop linker^72^. Both domains have their own paralogs with either stand-alone NTD or CTD; the former is called helical carotenoid protein (HCP) and the latter CTD-like proteins (CTDHs)^72^. *Anthocerobacter* has one OCP and one NTD, but no CTDH. While Grettenberger et al^12^ did report on the presence of OCP in *Candidatus* Aurora, they did not distinguish OCP, HCP, and CTDH. Our reanalysis showed that *Candidatus* Aurora has one OCP, one HCP, and one putative CTDH. We then incorporated the OCP sequences from *Anthocerobacter* and *Candidatus* Aurora into the dataset compiled by Bao *et al*^33^, which was used to define the major OCP clades^33^. Sequence alignment was done by PASTA v3^67^ followed by phylogenetic inference using IQTREE v2.0.3 with automatic model selection and 1,000 ultrafast bootstrap replicates^68^.

### Phylogeny of phycobilisome linker proteins

The phycobilisome linkers from *Anthocerobacter* and *Gloeobacter violaceus* were added to the protein sequences compiled by Watanabe and Ikeuchi^38^. Sequence alignment was done by PASTA v3^67^, and phylogeny was reconstructed using IQTREE v2.0.3 with automatic model selection and 1,000 ultrafast bootstrap replicates^68^.

### Microscopy

Pure cultures were processed for transmission electron microscopy as described previously^73^ with slight modifications. Cultures were fixed in 3% glutaraldehyde, 1% fresh formaldehyde, and 0.75 % tannic acid in 0.05 M Na-cacodylate buffer, pH 7, for 3 h at room temperature. After several rinses in 0.1M buffer, the samples were post-fixed in buffered (0.1M, pH 6.8) 1% osmium tetroxide overnight at 4°C, dehydrated in an ethanol series and embedded in Spurr’s resin via ethanol. Thin sections were cut with a diamond histo-knife, stained with methanolic uranyl acetate for 15 min and in Reynolds’ lead citrate for 10 min, and observed with a Hitachi H-7100 transmission electron microscope at the Imaging-Microscopy Platform of the IBIS, Universite Laval. Images of colonies on an agar plate were taken by a SMZ-171TLED stereomicroscope (Bio Pioneer Tech Co, Taiwan) equipped with a TrueChrome AF digital camera (TUCSEN, China). Light microscopy images were taken under a Leica DMR microscope (Leica, Germany) equipped with a Leica MC 170HD digital camera (Leica, Germany).

### Absorption and fluorescence spectral analyses

The spectroscopy measurements were performed as previously described with modifications^74^. The OD750 values of the cyanobacterial cultures were adjusted to 0.25 in 1 mL of B-HEPES medium for absorption spectroscopy, and 0.25 in 1 mL of PEG buffer (500 ul of B-HEPES medium and 500 ul of 30% (w/v) PEG4000) for fluorescence spectroscopy. The measurement of absorption spectra (wavelength from 400 to 800 nm) was performed by MAPADA UV1800 with an equipped PC software (Datech Instruments Ltd., Taiwan). Fluorescence spectroscopy was performed by the F-4500 fluorescence spectrophotometer (Hitachi, Japan). Fluorescence emission was recorded from 600 to 800 nm by excitation at 440 nm for chlorophyll *a*.

### Measurements of pigment composition and oxygen evolution

The quantifications of chlorophyll *a*, carotenoids, and phycobiliproteins were performed as previously described^75^. Steady-state rates of oxygen evolution were measured as previously described with minor modifications ^76^. An Oxytherm System oxygen electrode (Model V-560, Hansatech Instruments Ltd, UK) was used for the measurement. Cyanobacterial cells were diluted in B-HEPES medium to the final concentration as 3.5 μg (Chl) ml^−1^ in a stirred, water-jacketed cell chamber at 25°C. Potassium ferricyanide and 2,6-dichloro-p-benzoquinone (DCBQ) were added sequentially into the cell chamber as artificial electron acceptors to the final concentration as 2 mM. Different strengths of light intensities were provided from both sides of the cell chamber by two fiber-optic illuminators (Dolan-Jenner model MI 150, USA). Data recording and processing were performed with the manufacture software O2 view (Hansatech Instruments Ltd, UK).

## Acknowledgments

We thank Dr. Noris Salazar Allen for providing fieldwork support to J.C.V.A., Dr. Hsiu-An Chu and Keng-Min Lin for assisting oxygen evolution measurement, Dr. Patrick Shih for sharing alignment matrices and early discussions, and Drs. Kathleen Pryer and John Meeks for comments. This work was funded by National Science Foundation (NSF) grant no. DEB1831428 to F.-W.L., Ministry of Science and Technology (Taiwan) Young Scholar Fellowship Einstein Program (108C5368 and 109C5369) and Ministry of Education (Taiwan) Yushan Young Scholar Program (108V1102 and 109V1102) to M.-Y.H, and Chaires de Recherche du Canada (950-232698), CRNSG (RGPIN-2016-05967) and Earl S. Tupper Fellowship (STRI) to J.C.V.A.

## Author Contributions

F.-W.L. and M.-Y.H. designed research; N.R., D.A.H., J.M.N., P.Y.C., J.C.V.A., M.-Y.H., F.-W.L. performed research; N.R., M.-Y.H., and F.-W.L. analyzed data; and N.R., M.-Y.H., and F.-W.L. wrote the paper.

