## Supplementary material for "Revisiting the early evolution of Cyanobacteria with a new thylakoid-less and deeply diverged isolate from a hornwort": Fig. S1-7

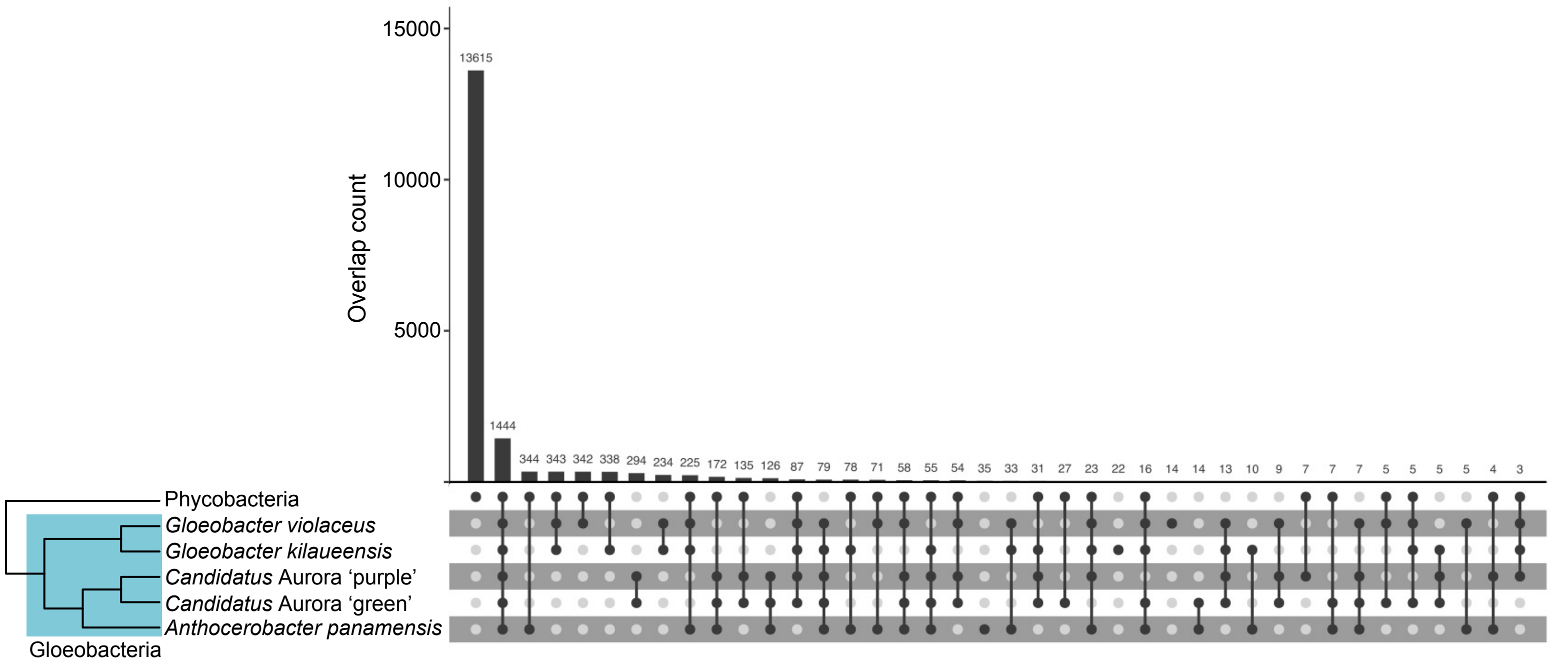


Figure S1. **Protein family overlaps among Gloeobacteria (5 species) and Phycobacteria (98 species) from Orthofinder.**

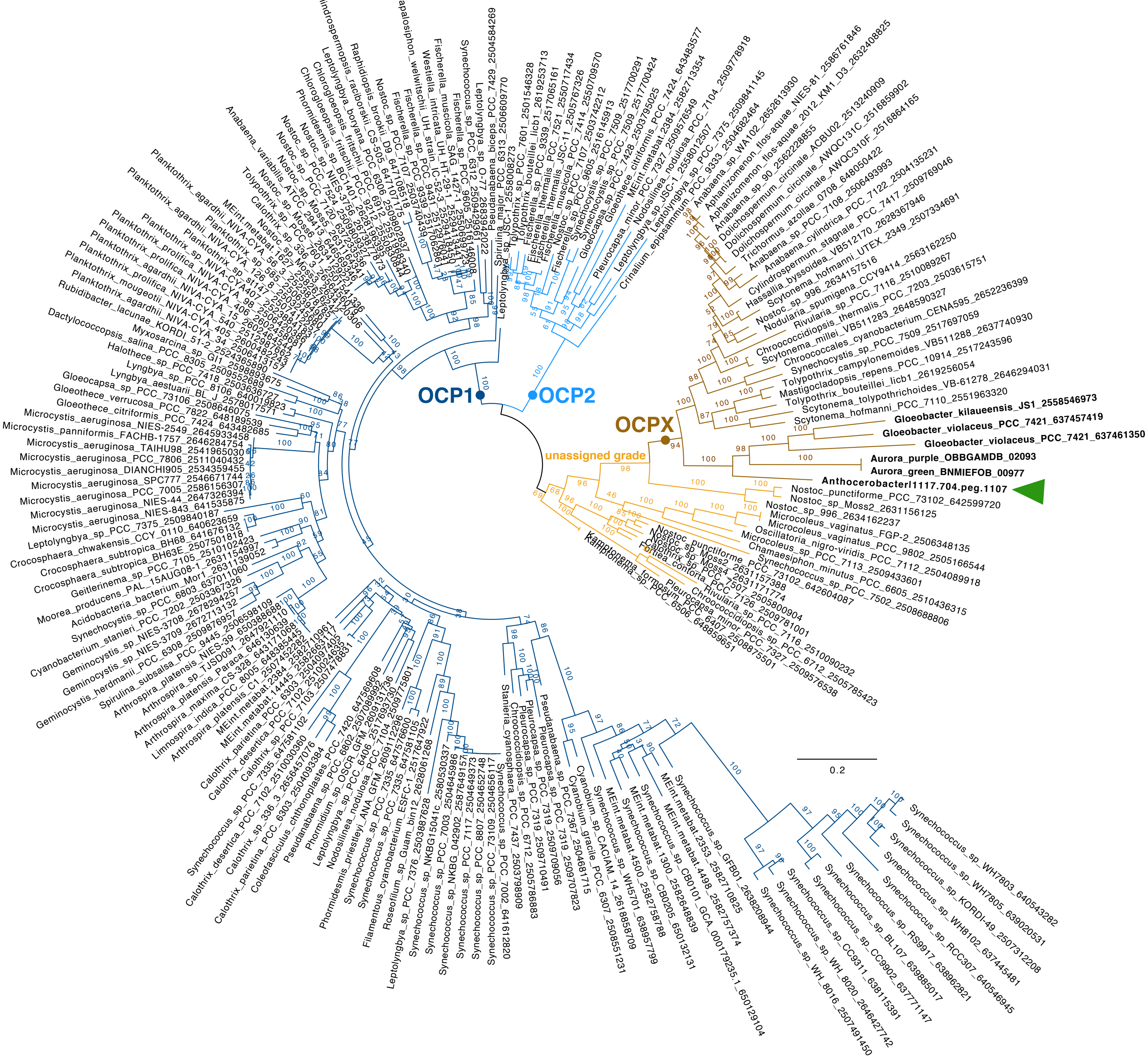


Figure S2. **Phylogeny of orange carotenoid proteins (OCP).** *Anthocerobacter* has one OCP homolog (green arrowhead) belonging to the OCPX clade. The clade designation followed that of Bao et al (2017). The numbers above branches are bootstrap support values.

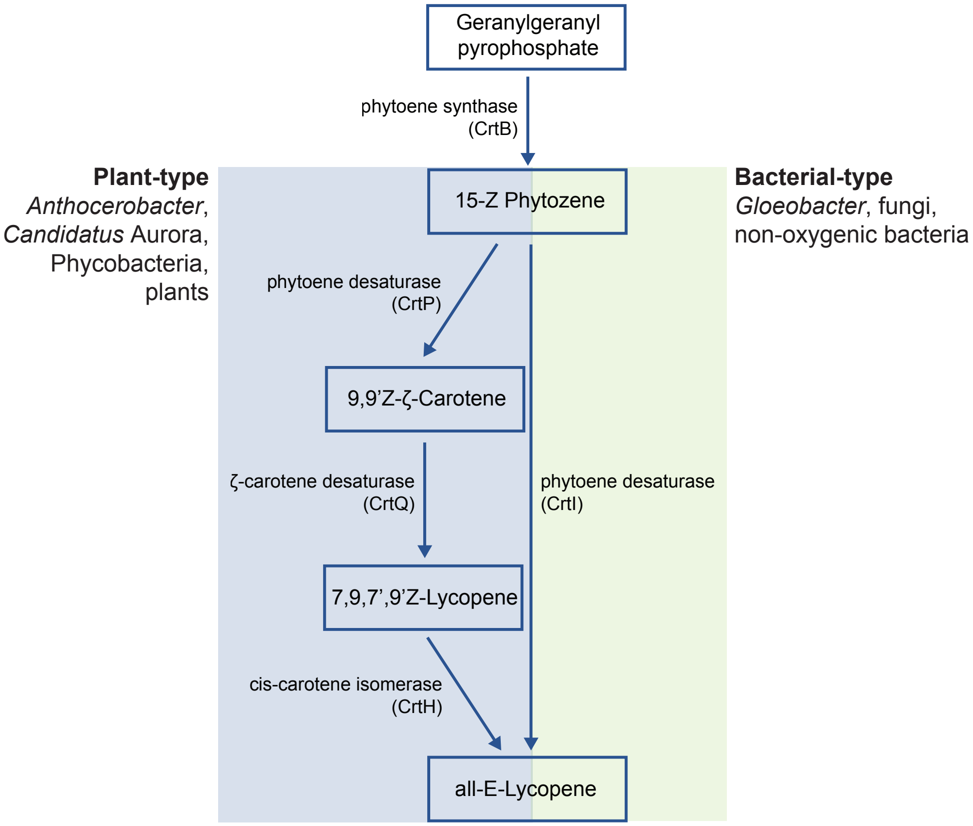


Figure S3. **Comparison of plant-type and bacterial-type carotenoid biosynthesis pathways.** *Anthocerobacter* has the plant-type pathway like most other Cyanobacteria, whereas its sister genus *Gloeobacter* has the bacterial-type pathway.

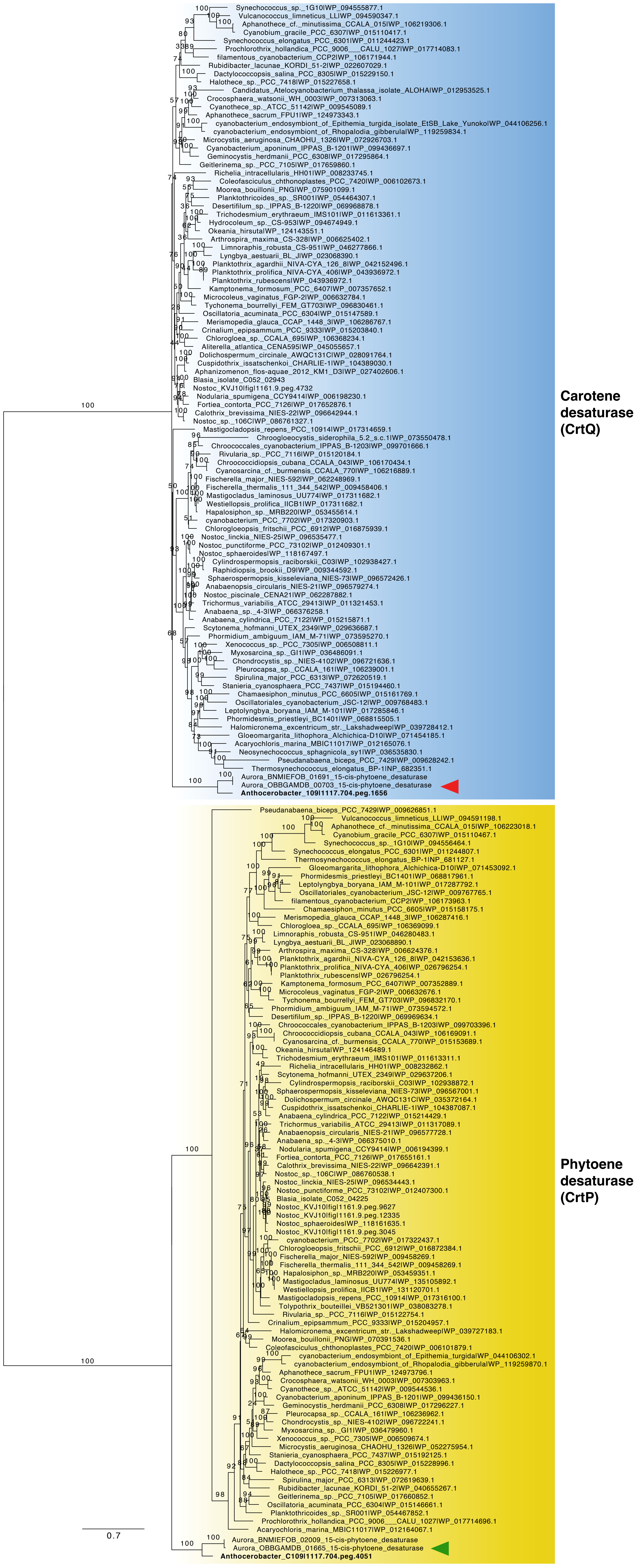


Figure S4. **Phylogeny of cyanobacterial CrtP and CrtQ (orthogroup OG0000256).** The *Anthocerobacter* and *Candidatus* Aurora CrtP orthologs (green arrowhead) are sister to Phycobacteria with strong support. For CrtQ, the orthologs (red arrowhead) are one-step nested within Phycobacteria sequences but without strong backbone support. These results do not support horizontal gene acquisition of *Anthocerobacter* CrtP and CrtQ from Phycobacteria, but instead imply that both genes were present in the most recent common ancestor of Cyanobacteria. The numbers above branches are bootstrap support values.

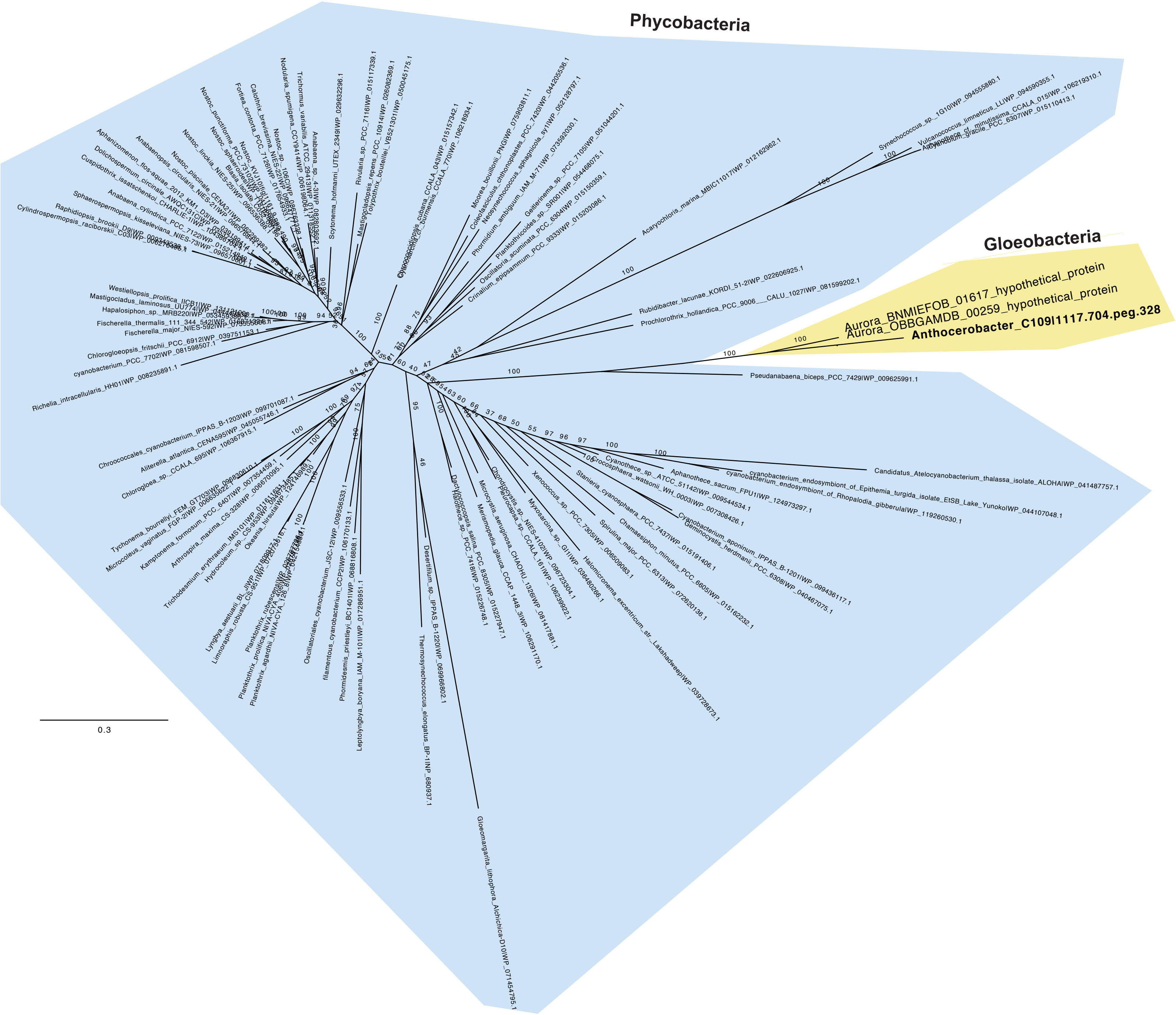

Figure S5. **Phylogeny of Carotene isomerase CrtH (orthogroup OG0001645).** The tree is not rooted because no close and alignable outgroup sequences can be found. Nevertheless, the *Anthocerobacter* and *Candidatus* Aurora orthologs (yellow background) are distinct from those of Phycobacteria (blue background), suggesting that recent horizontal gene transfer is unlikely. The numbers above branches are bootstrap support values.

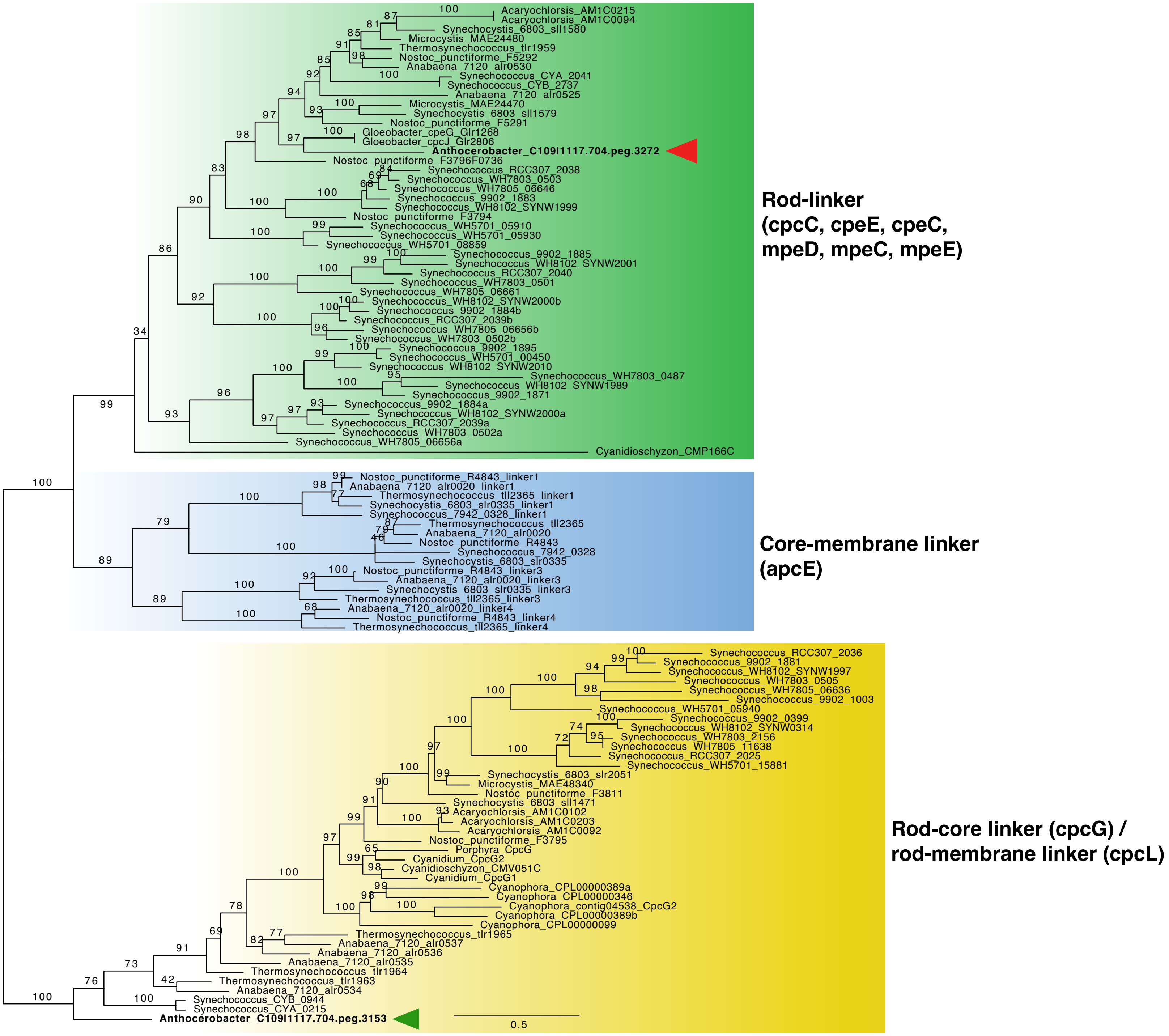


Figure S6. **Phylogeny of phycobilisome linkers.** Unlike *Gloeobacter* and *Candidatus* Aurora, *Anthocerobacter* has a cpcG homolog (green arrowhead). The *Anthocerobacter* cpcG is sister to the rest of cyanobacterial cpcG or cpcL copies with strong bootstrap support, suggesting that horizontal gene transfer is unlikely and that cpcG is probably present in the most common recent ancestor of Cyanobacteria. The numbers above branches are bootstrap support values. The *Anthocerobacter* cpcJ homolog is marked by a red arrowhead.

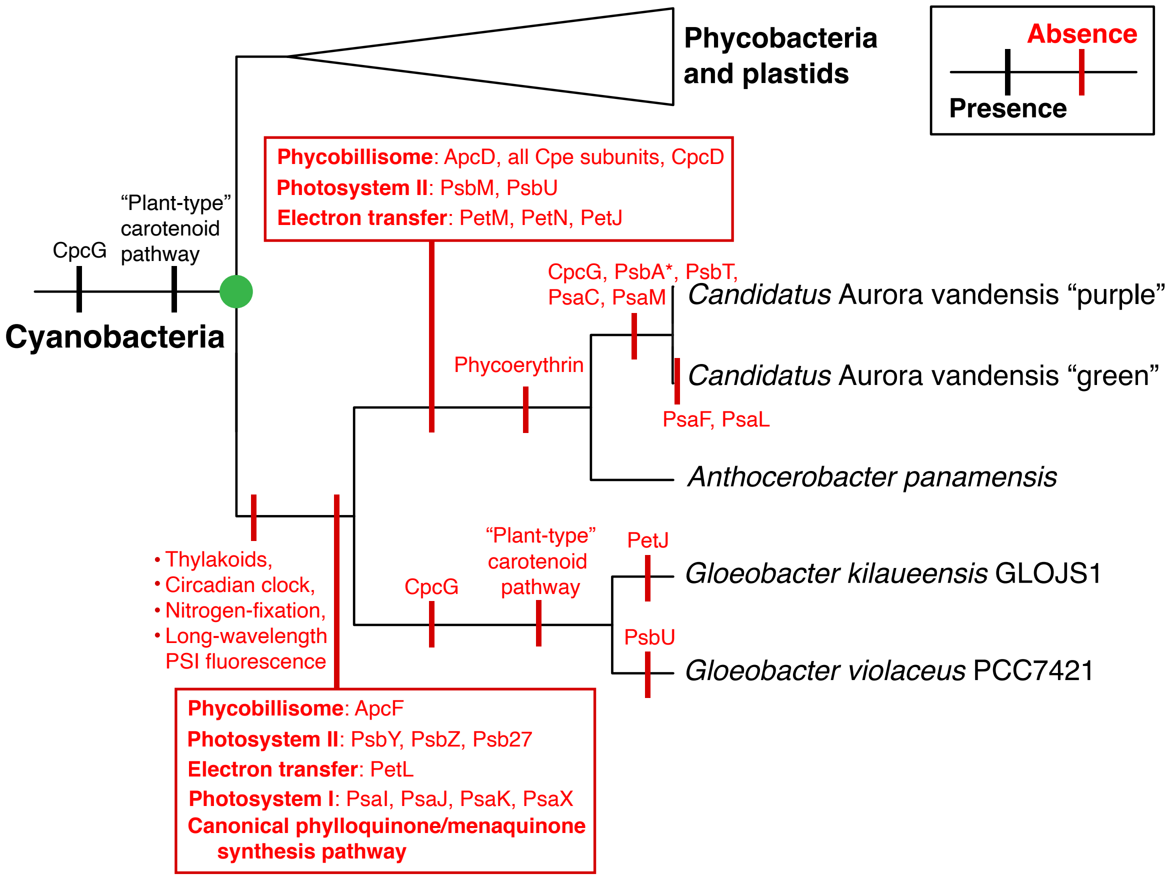


Figure S7. **Synapomorphies of different Gloeobacteria lineages.** Shared presence/absence of selected traits or protein components is mapped along the branches. *psbA was not found in *Candidatus* Aurora metagenome assembly, but on a separate contig that was not properly binned.
